# The eccDNA mediated spread and rapid evolution of herbicide resistance in pigweed interspecific hybrids

**DOI:** 10.1101/2023.03.01.530670

**Authors:** Dal-Hoe Koo, Rajendran Sathishraj, Sridevi Nakka, Yoonha Ju, Vijay K. Nandula, Mithila Jugulam, Bernd Friebe, Bikram S. Gill

**Author notes:** **Corresponding Author**: Dal-Hoe Koo.

## Abstract

Extrachromosomal circular DNAs (eccDNAs) are found in many eukaryotic organisms. The eccDNA-powered copy number variation plays diverse roles from oncogenesis in humans to herbicide resistance in crop weeds. Here we report interspecific eccDNA flow and its dynamic behavior in soma cells of natural populations and F_1_ hybrids of *Amaranthus* sp. The glyphosate resistance (GR) trait is controlled by eccDNA-based amplification harboring the *5-enolpyruvylshikimate-3-phosphate synthase* (*EPSPS*) gene (*EPSPS*-eccDNA), the molecular target of glyphosate. We documented pollen-mediated transfer of eccDNA in experimental hybrids between GS *A. tuberculatus* x GR *A. palmeri*. Experimental hybridization and fluorescence *in situ* hybridization (FISH) analysis revealed that the *EPSPS*-eccDNA present in *A. spinosus* was derived from GR *A. palmeri* by natural hybridization. FISH analysis also revealed random chromosome anchoring and massive *EPSPS*-eccDNA copy number variation in soma cells of weedy hybrids. The results suggest that eccDNAs are inheritable across compatible species contributing to genome plasticity and rapid adaptive evolution.

## Introduction

Extrachromosomal circular DNA (eccDNA) is generated from chromosomal DNAs in sizes ranging from a few hundred base pairs up to megabases, and they contribute to genome plasticity in eukaryotes (Stark and Wahl, 1984; Cohen et al., 2008; Pennisi, 2017; Koo et al., 2018). High copy number and expression of eccDNA drives the oncogenesis and promotes the survival and proliferation of cancerous cells in humans (Turner et al., 2017). In plants, Koo et al. (2018) reported that glyphosate resistance in a troublesome crop weed *Amaranthus palmeri* was driven by eccDNA, which contain the 5-enolpyruvylshikimate-3-phosphate synthase (*EPSPS*) gene the molecular target of glyphosate (Steinrücken and Amrhein, 1980). This specific eccDNA is, hereafter referred as *EPSPS*-eccDNA. The glyphosate resistance may involve target site alterations, either by mutation or amplification of the *EPSPS* gene (Gaines et al., 2010; Dillon et al., 2017). The amplification of the *EPSPS* gene conferring glyphosate resistance in *A. palmeri* was first documented in Georgia, in 2006 and has since spread to most states in the US. All analyzed glyphosate-resistant (GR) populations carry the identical *EPSPS*-eccDNA (Koo et al., 2018; Molin et al., 2018). Nandula et al. (2014) reported the apparent transfer of glyphosate resistance from *A. palmer* to related species, *A. spinosus* L., another troublesome weed. The analysis of GR *A. spinosus* genotypes showed amplification of *EPSPS* gene (up to 40 copies) and their gene sequence was identical to the *EPSPS* gene of GR *A. palmeri* (Nandula et al., 2014). This led to the hypothesis that glyphosate resistance found in *A. spinosus* arose from pollen-mediated *EPSPS*-eccDNA transfer from GR *A. palmeri*. Our results confirm this hypothesis. Furthermore, we present results on the dynamics of *EPSPS*-eccDNA driven evolution of herbicide resistance in an interspecific hybrid between *A. tuberculatus* (female, 2n=32) and GR *A. palmeri* (male, 2n=34).

## Results

### Pollen-mediated *EPSPS*-eccDNA flow between species

Fiber-FISH analysis using six BACs [used in Koo et al. (2018)] associated with *EPSPS*-eccDNA of GR *A. palmeri* detected two types of eccDNAs in GR *A. spinosus* (**Fig. 1**), circular and linear with structural polymorphisms similar to those found in GR *A. palmeri* (Koo et al., 2018). The results indicated transfer of *EPSPS*-eccDNA from GR *A. palmeri* to GR *A. spinosus via* pollen by interspecific hybridization in the natural environment. To test further the pollen-mediated *EPSPS*-eccDNA transfer by interspecific inheritance, six interspecific F_1_ hybrids were generated from a cross between *A. tuberculatus* (female, 2n=32) and GR *A. palmeri* (male, 2n=34) carrying *EPSPS*-eccDNA. Molecular analysis of six F_1_ hybrids using species-specific markers revealed that two F_1_ hybrids (W-P#3 and W-P#6) were positive for both the markers tested while the other four F_1_ hybrids were negative for *A. palmeri* specific marker (**Fig. 2*a***), whereas all the six F_1_ hybrids were positive for *A. tuberculatus* specific marker (**Fig. 2*b***).

**Figure 1.**
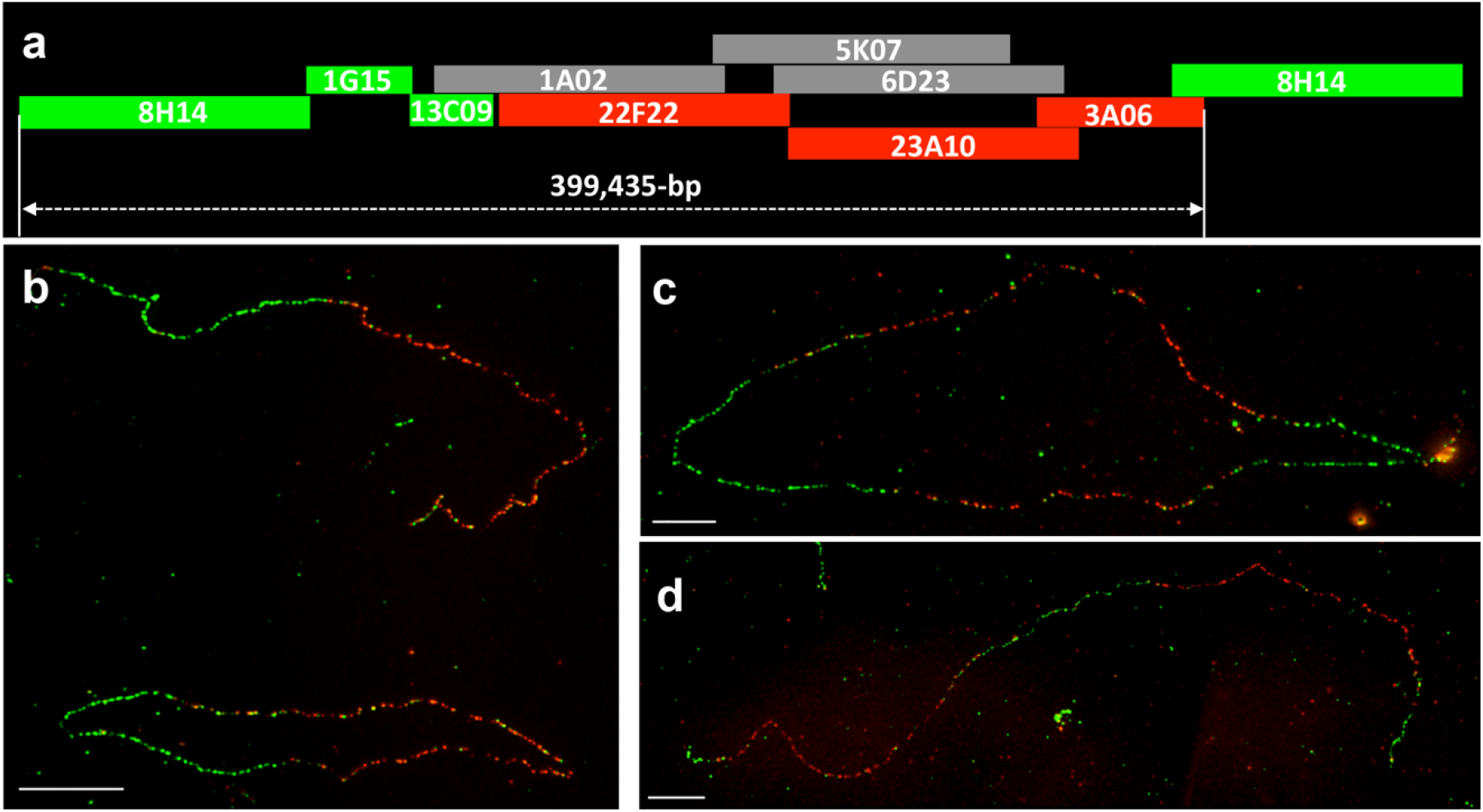
Fiber-FISH images of *EPSPS*-eccDNAs in GR *A. spinosus*. **a**, schematic representation of BAC-contig assembly of eccDNA used in fiber-FISH analysis (Koo et al., 2018); **b,** a circular form (bottom) and linear form of eccDNA (top); **c**, dimerized circular form of eccDNA with head-to-tail tandem orientation; **d**, linear form of eccDNA with head-to-tail dimer. Green colors: BAC 8H14, BAC 1G15, BAC 13C09. Red colors: BAC 22F22, BAC 23A10, BAC 3A06. Bars=10 μm

### Random chromosome tethering and somatic mosaicism for copy number variation of *EPSPS*-eccDNA in F_1_ hybrids

FISH analysis detected five pairs of chromosomes carrying the 5S rDNA signals in *A. tuberculatus* (**Fig. 2*c***) and one pair in *A. palmeri* (**Fig. 2*d***). As expected, the 5S rDNA-FISH detected six major hybridization signals in the metaphase chromosomes of F_1_ hybrid, W-P#3; five from *A. tuberculatus* and one from *A. palmeri* (**Figs. 2*e-g***). The 5S rDNA-FISH marker chromosomes provided an opportunity to study the dynamics of *EPSPS*-eccDNA anchoring to chromosomes of interspecific origin. The *EPSPS*-eccDNA FISH signals were randomly associated with 5S rDNA-marker chromosome of *A. tuberculatus* in the F_1_ hybrid plant, W-P#3. One cell had eccDNA tethered with four of the five chromosomes of *A. tuberculatus* and with one chromosome of *A. palmeri* (**Fig. 2*e***). The second cell had eccDNA tethered with only one of the five chromosomes of *A. tuberculatus* and with one chromosome of *A. palmeri* (**Fig. 2*f***). The third cell had no tethering of eccDNA with any of the 5S rDNA-marker chromosomes (**Fig. 2*g***). The analysis provided conclusive evidence that the *EPSPS*-eccDNA of *A. palmeri* is capable of random tethering to chromosomes of a related species.

**Figure 2.**
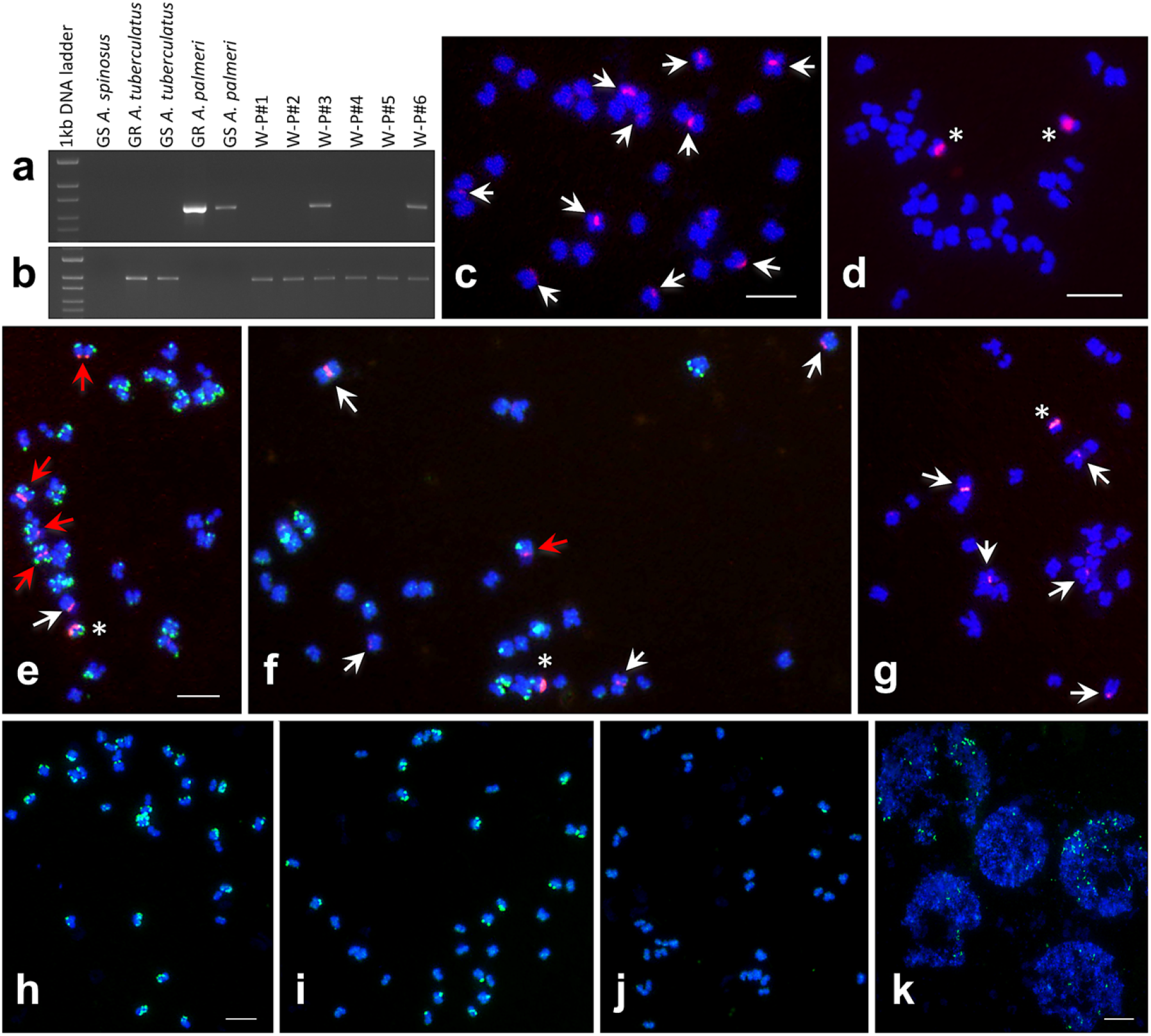
Conformation of F_1_ hybrids by PCR-based amplification and characterization of F_1_s along with their parents by FISH. Amplification of *A. palmeri* specific 697 bp genomic region using AW155/AW90 markers (**a**). Amplification of *A. tuberculatus* specific 992 bp genomic region using AW468/AW469 markers (**b**). Two F_1_ hybrids (W-P#3 and W-P#6) were positive for both the markers tested. FISH mapping of 5S rDNAs on mitotic metaphase chromosomes of *A. tuberculatus* (2n=32) (white arrows in **c**) and *A. palmeri* (2n=34) (white asterisks in **d**). Distribution of eccDNAs (green signals) on the 5S rDNA-marker chromosomes of *A. tuberculatus* in F_1_ hybrid from single root tip (**e-g**). Out of five 5S rDNA-marker chromosomes of *A. tuberculatus* (arrows) four (red arrows), (**e**) one (**f**), and none (**g**) of them had eccDNAs. The 5S rDNA-marker chromosome from *A. palmeri* is marked by an asterisk. Distribution of eccDNAs (green signals) on the chromosomes of GR *A. spinosus* from single root tip (**h-k**). *EPSPS*-eccDNAs (green signals) associated with most (**h**), half (**i**), and few (**j**) of the mitotic metaphase chromosomes of GR *A. spinosus* and mosaic pattern of *EPSPS*-eccDNA copy numbers in interphase cells of the same tissue (**k**). Bars=5 μm.

Apart from random tethering, somatic mosaicism for copy number variation of *EPSPS*-eccDNA was observed in GR *A. spinosus* and F_1_ hybrids. In a sample of three cells of GR *A. spinosus* (**Figs. 2*h-j***), one cell had *EPSPS*-eccDNA signals on most of the chromosomes (**Fig. 2*h***; pattern I), the second cell had *EPSPS*-eccDNA signals on half of the chromosomes (**Fig. 2*i***; pattern II) and the third cell had few *EPSPS*-eccDNA signals (**Fig. 2*j***; pattern III). The frequency of patterns I, II, III observed in a sample of 50 cells of GR *A. spinosus* having *EPSPS* gene copy number of 54.3 ± 2.3 (**Table 1**) was 30%, 54% and 2%, respectively. In F_1_ hybrid plant W-P#3 (**Figs. 2*f-g***) (n=30) where the *EPSPS* gene copy number was 3.0 ± 0.8 (**Table 2**); the observed frequency of *EPSPS*-eccDNA signals corresponding to patterns I, II and III was 20%, 66.7% and 13%, respectively. Similar cytological behavior of *EPSPS*-eccDNA was also observed in the W-P#6 genotype (**Table 2**).

**Table 1.** *EPSPS* gene and *EPSPS*-eccDNA copy number in glyphosate-susceptible (GS) and – resistant (GR) *A. spinosus*, and GR *A. palmeri* plants. The relative *EPSPS:β-tubulin* gene copy number was adjusted to 1 for known GS *A. spinosus* and the copy numbers for GR *A. palmeri and A. spinosus* plants shown here are relative to the GS *A. spinosus*

| Plant | *EPSPS* copy number<br>(No. ± SD) | *EPSPS*-eccDNA<br>(BAC-FISH) |
| --- | --- | --- |
| GS *A. spinosus* | 1.0±0.9 | 0% (n=100) |
| GR *A. palmeri* | 77.3±8.6 | >90% (Koo et al., 2018) |
| GR *A. spinosus*-1* | 127.5±4.7 | fiber-FISH |
| GR *A. spinosus*-2 | 97.4±10.7 | nd |
| GR *A. spinosus*-4 | 126.6±7.1 | nd |
| GR *A. spinosus*-5 | 54.3±2.3 | 98% (n=50) |
| GR *A. spinosus*-7 | 62.4±8.5 | nd |
| GR *A. spinosus*-8 | 46.3±1.4 | nd |

Asterisk indicates the plant that used in fiber-FISH analysis
%: mitotic metaphase cell with BAC-FISH signal.
nd: not determined.

**Table 2.** *EPSPS* gene and eccDNA copy number in interspecific F_1_ hybrids *A. tuberculatus* (1.0±0.1) x *A. palmeri* (77.3±8.6). The relative *EPSPS:β-tubulin* gene copy number was adjusted to 1 for known GS *A. palmeri* and the copy numbers for interspecific hybrids shown here are relative to the GS *A. palmeri*

| Plant | *EPSPS* copy number<br>(No. ± SD) | *EPSPS*-eccDNA (BAC-FISH) |
| --- | --- | --- |
| GS *A. palmeri* | 1.0 ± 0.1 | 0% (Koo et al., 2018) |
| W-P#3 | 3.0 ± 0.8 | 87% (n=30) |
| W-P#6 | 4.4 ± 0.5 | 93.3% (n=30) |

%: mitotic metaphase cell having BAC-FISH signal.

## Discussion

Nandula et al. (2014) reported the first case of GR *A. spinosus* as a result of amplification of the *EPSPS* gene. The differential homology of *EPSPS* gene sequence between GR and GS *A. spinosus* with that of GR *A. palmeri* hypothesized that glyphosate resistance in *A. spinosus* was due to pollen-mediated transfer of glyphosate resistant gene by hybridization with naturally occurring population of GR *A. palmeri*. Fiber-FISH and FISH analysis of GR *A. spinosus* supported this hypothesis and proved that GR *A. spinosus* harbors *EPSPS*-eccDNA (**Fig. 1** and **Fig. 2**), which it most likely acquired from GR *A. palmeri* through interspecific hybridization as both species are naturally compatible to hybridization (Franssen et al., 2001). Further, controlled hybridization experiment was conducted to demonstrate the transferability of *EPSPS*-eccDNA during interspecific hybridization. Six F_1_ hybrids were developed by hybridizing *A. palmeri* with *A. tuberculatus*. Cytological and molecular analyses of F_1_ hybrids corroborated that *EPSPS*-eccDNA from *A. palmeri* can be transferred by interspecific hybridization (**Fig. 2**). *A. palmeri, A. tuberculatus* and *A. spinosus* are members of family Amaranthaceae and are among the most problematic weeds in the US. Their floral biology and close genetic relationships favor interspecific hybridization among these species and enable the spread of herbicide resistance (Oliveira et al., 2018). Earlier we reported the *EPSPS*-eccDNA mediated evolution of glyphosate resistance in *A. palmeri* (Koo et al., 2018). This report documents *A. palmeri EPSPS*-eccDNA is capable of pollen-mediated transfer and spread to related species posing a serious burden on control of pigweeds in crops.

Hybridization of GR *A. palmeri* having *EPSPS* copy number of 77.3 ± 8.6 with GS *A. tuberculatus* having *EPSPS* copy number of 1 resulted in true F_1_ hybrid W-P#3 with *EPSPS* copy number of 3.0 ± 0.8 and was susceptible to glyphosate (**Table 2**). The FISH experiments revealed that W-P#3 has a certain percentage of soma cells in its meristematic tissues that harbor a large number of *EPSPS*-eccDNA signals anchored to the chromosomes (**Figs. 2*f-g***). The cells with a large number of *EPSPS*-eccDNA copies can potentially survive glyphosate application, multiply and produce resistant shoots during the lifetime of this hybrid. Some of the seed borne on the resistant shoots will transmit the glyphosate resistance to the progeny. In cell culture studies exposed to drugs, similar kind of resistance mediated by episomes (here referred as eccDNA) has been described as an acquired resistance or trait (Stark and Wahl, 1984). Our work on *A. palmeri* (Koo et al., 2018) and this report are documented examples of rapid adaptive evolution of a trait in higher organisms mediated by eccDNA. The eccDNAs enable copy number variation in soma cells during the growth of the organism which under strong selection pressure may drive rapid adaptive evolution, and support the Lamarckian theory of acquired characters [see also Koo et al. (2018)].

## Materials and Methods

### Plant materials

*A. spinosus* plants were obtained from the field in Lafayette County, Mississippi (34.23450 N, 89.63433 W), and were confirmed to be glyphosate resistant (Nandula et al., 2014). The collected plants were cloned by vegetative propagation in the greenhouse. The interspecific hybrids between *A. tuberculatus* and *A. palmeri* were produced as per the procedure given by Gaines et al. (Gaines et al., 2012). A total of 6 interspecific F_1_ plants were randomly selected for FISH analysis of eccDNAs.

### Molecular analysis

The quantitative PCR procedure given by Koo et al. (Koo et al., 2018) was followed to determine the *EPSPS* copy number in GR *Amaranthus* sp. and their interspecific hybrids. PCR based species-specific primers developed by Wright et al. (Wright et al., 2016) were used for parental species screening and evaluation of F_1_ hybrids.

### Fluorescence *in situ* hybridization (FISH)

The procedures for mitotic chromosome preparation, FISH and fiber-FISH were adapted from Koo et al. (2018). The images were captured with a Zeiss Axioplan 2 microscope (Carl Zeiss Microscopy LLC, Thornwood, NY) using a cooled CCD camera CoolSNAP HQ2 (Photometrics, Tucson, AZ) and AxioVision 4.8 software. The final contrast of the images was processed using Adobe Photoshop CS5 software.

## Acknowledgments and Funding

We thank W. John Raupp for critical review of the manuscript and Duane Wilson for technical assistance. This research was supported by grants from the Kansas Wheat Commission and the Kansas Crop Improvement Association, Wheat Genetics Resource Center Industry/ University Cooperative Research Center National Science Foundation Contract 1822162. This is contribution number 23-216-J from the Kansas Agricultural Experiment Station, Kansas State University, Manhattan, KS 66506-5502, USA.

## Author Contributions

D-HK, RS, SN, and YJ performed the experiments; D-HK and BSG wrote the manuscript; VKN confirmed and collected the glyphosate-resistant *A. spinosus;* D-HK, BF, and BSG designed the experiments; D-HK, RS, YJ, SN, VKN, MJ, BF, and BSG analyzed the data and helped to draft the final manuscript.

## Parsed Citations

**Cohen S, Houben A, Segal D (2008) Extrachromosomal circular DNA derived from tandemly repeated genomic sequences in plants. The Plant Journal 53: 1027-1034**

Google Scholar: <u>Author Only Title Only Author and Title</u>

**Dillon A, Varanasi VK, Danilova TV, Koo D-H, Nakka S, Peterson DE, Tranel PJ, Friebe B, Gill BS, Jugulam M (2017) Physical mapping of amplified copies of the 5-enolpyruvylshikimate-3-phosphate synthase gene in glyphosate-resistant Amaranthus tuberculatus. Plant physiology 173: 1226-1234**

Google Scholar: <u>Author Only Title Only Author and Title</u>

**Franssen AS, Skinner DZ, Al-Khatib K, Horak MJ, Kulakow PA (2001) Interspecific hybridization and gene flow of ALS resistance in Amaranthus species. Weed Science 49: 598-606**

Google Scholar: <u>Author Only Title Only Author and Title</u>

**Gaines TA, Ward SM, Bukun B, Preston C, Leach JE, Westra P (2012) Interspecific hybridization transfers a previously unknown glyphosate resistance mechanism in Amaranthus species. Evolutionary applications 5: 29-38**

Google Scholar: <u>Author Only Title Only Author and Title</u>

**Gaines TA, Zhang W, Wang D, Bukun B, Chisholm ST, Shaner DL, Nissen SJ, Patzoldt WL, Tranel PJ, Culpepper AS (2010) Gene amplification confers glyphosate resistance in Amaranthus palmeri. Proceedings of the National Academy of Sciences 107: 10291034**

Google Scholar: <u>Author Only Title Only Author and Title</u>

**Koo D-H, Molin WT, Saski CA, Jiang J, Putta K, Jugulam M, Friebe B, Gill BS (2018) Extrachromosomal circular DNA·based amplification and transmission of herbicide resistance in crop weed Amaranthus palmeri. Proceedings of the National Academy of Sciences 115: 3332-3337**

Google Scholar: <u>Author Only Title Only Author and Title</u>

**Molin WT, Wright AA, VanGessel MJ, McCloskey WB, Jugulam M, Hoagland RE (2018) Survey of the genomic landscape surrounding the 5-enolpyruvylshikimate-3-phosphate synthase (EPSPS) gene in glyphosate-resistant Amaranthus palmeri from geographically distant populations in the USA. Pest management science 74: 1109-1117**

Google Scholar: <u>Author Only Title Only Author and Title</u>

**Nandula VK, Wright AA, Bond JA, Ray JD, Eubank TW, Molin WT (2014) EPSPS amplification in glyphosate-resistant spiny amaranth (Amaranthus spinosus): a case of gene transfer via interspecific hybridization from glyphosate-resistant Palmer amaranth (Amaranthus palmeri). Pest management science 70: 1902-1909**

Google Scholar: <u>Author Only Title Only Author and Title</u>

**Oliveira MC, Gaines TA, Patterson EL, Jhala AJ, Irmak S, Amundsen K, Knezevic SZ (2018) Interspecific and intraspecific transference of metabolism-based mesotrione resistance in dioecious weedy Amaranthus. The Plant Journal 96: 1051-1063**

Google Scholar: <u>Author Only Title Only Author and Title</u>

**Pennisi E (2017) Circular DNA throws biologists for a loop. Science 356: 996-996**

Google Scholar: <u>Author Only Title Only Author and Title</u>

**Stark GR, Wahl GM (1984) Gene amplification. Annual review of biochemistry 53: 447-491**

Google Scholar: <u>Author Only Title Only Author and Title</u>

**Steinrücken H, Amrhein N (1980) The herbicide glyphosate is a potent inhibitor of 5-enolpyruvylshikimic acid-3-phosphate synthase. Biochemical and biophysical research communications 94: 1207-1212**

Google Scholar: <u>Author Only Title Only Author and Title</u>

**T urner KM, Deshpande V, Beyter D, Koga T, Rusert J, Lee C, Li B, Arden K, Ren B, Nathanson DA (2017) Extrachromosomal oncogene amplification drives tumour evolution and genetic heterogeneity. Nature 543: 122-125**

Google Scholar: <u>Author Only Title Only Author and Title</u>

**Wright AA, Molin WT, Nandula VK (2016) Distinguishing between weedy Amaranthus species based on intron 1 sequences from the 5-enolpyruvylshikimate-3-phosphate synthase gene. Pest management science 72: 2347-2354**

Google Scholar: <u>Author Only Title Only Author and Title</u>

